# Beyond the urban surface: bird-borne sensing reveals the atmospheric structure of urban heat

**DOI:** 10.64898/2026.10.12.758485

**Authors:** Giacomo Grosso, Claudio Carere, Giacomo Dell’Omo

## Abstract

Urban heat is commonly characterized near ground level, while its vertical atmospheric structure remains difficult to observe across cities. We used bird-borne temperature sensors on homing pigeons (*Columba livia*) and yellow-legged gulls (*Larus michahellis*) to investigate the horizontal, vertical and daytime structure of the lower urban atmosphere over Rome. The dataset comprised 61,917 air-temperature observations from 47 individuals of gulls and pigeons, spanning 17 km and altitudes up to 400 m above ground level. Generalized additive models related air temperature to distance from the city centre, altitude, time of day, land cover, tree-patch area and topography, and reconstructed three-dimensional temperature fields. Air temperature showed a centre-to-periphery gradient that weakened with altitude and through the daytime period. At 20 m above ground level, its magnitude declined from −0.40 °C km⁻¹ at 09:00 to −0.30 °C km⁻¹ at 13:00 and −0.19 °C km⁻¹ at 17:00, with weaker gradients at higher altitudes. Reconstructed isothermal boundaries showed a complementary vertical pattern. At 13:00, median heights decreased from 135 m for the 34 °C boundary to 76 m for 35 °C and 37 m for 36 °C. The 34 °C boundary reached its greatest extent around midday, rising from 57 m at 09:00 to 135 m at 13:00 before declining to 14 m at 17:00. Horizontal thermal contrast and the vertical development of high-temperature air represent complementary dimensions of urban heat. Bird-borne sensing offers a promising approach for resolving three-dimensional atmospheric temperature patterns that conventional monitoring networks cannot capture.

## 1. Introduction

Urbanization is one of the most widespread forms of land transformation worldwide and increasingly interacts with climate change to alter local and regional climatic conditions (Grimmond 2007; McCarthy et al. 2010; IPCC 2023). One of its clearest effects is the urban heat island (UHI), broadly defined as higher temperatures in urban areas than in their less urbanized surroundings (Oke 1982; Arnfield 2003; Santamouris 2015). Urban warming results from changes in land cover, surface geometry and surface–atmosphere energy exchanges. These include the replacement of vegetated and permeable surfaces by impervious materials, reduced evapotranspiration, increased heat storage, radiative trapping within the urban canopy and anthropogenic heat release (Oke 1982; Voogt and Oke 2003; Gago et al. 2013). These changes can increase energy demand and human exposure to heat, with important consequences for health, especially during summer and heatwave events (Santamouris 2015; Heaviside et al. 2017).

Urban heat islands can be studied using either surface or air temperature. The surface urban heat island (SUHI) is generally described from land surface temperature, commonly obtained by thermal remote sensing, whereas the atmospheric urban heat island (AUHI) is based on air-temperature differences between urban and less urbanized areas (Voogt and Oke 1997, 2003; Wang and Li 2021). These two measures are related but do not describe the same thermal conditions and may show different spatial and temporal patterns. Air temperature varies considerably within a city. Land cover, vegetation, urban morphology, topography and meteorological conditions can all contribute to this variability (Oke 1987; Arnfield 2003; Ramamurthy and Bou-Zeid 2017).

Urban effects on air temperature are not restricted to the near-surface layer. They can extend into the urban boundary layer (UBL), the part of the atmospheric boundary layer influenced by the underlying urban surface (Oke 1987; Arnfield 2003). During daytime, surface heating and convective mixing redistribute heat vertically, while local advection also contributes to its horizontal transport. Consequently, the temperature difference between different parts of a city can change with both altitude and time of day (Miao et al. 2009; Hidalgo et al. 2010; Ramamurthy and Bou-Zeid 2017). Under weak background winds, differences in temperature between urban and surrounding areas may also generate an urban-breeze circulation. In idealized conditions, this circulation is characterized by low-level convergence towards the warmer urban area, upward motion over the city and divergence at higher levels, producing a dome-like structure (Fan et al. 2017). Modern cities are more complex. Topography, urban morphology, atmospheric stability and background winds can alter this structure, and sufficiently strong winds may displace or deform it into an asymmetric or plume-like pattern (Fan et al. 2017; Zhang et al. 2025). A dome-like temperature pattern alone, however, does not demonstrate the presence of a full heat-dome circulation, which would require measurements of wind fields, vertical motion and boundary-layer dynamics.

Observing this spatial and vertical variability directly remains difficult at the scale of a city. Fixed meteorological stations provide detailed time series but are often too sparse to capture intra-urban spatial variability, while mobile ground-based transects mainly sample the near-surface atmosphere (Oke 2006; Muller et al. 2013). Thermal remote sensing provides extensive spatial coverage but measures land surface rather than air temperature (Voogt and Oke 2003; Weng et al. 2004; Liu and Weng 2008). Numerical models can reconstruct atmospheric conditions in three dimensions but require observational data for evaluation (Masson 2006; Ramamurthy and Bou-Zeid 2017). Direct vertical observations from towers, radiosondes, aircraft or atmospheric profilers are generally restricted to a limited number of locations or specific campaigns. Repeated observations combining broad horizontal coverage with vertical sampling of urban air temperature therefore remain uncommon at the metropolitan scale.

Animal-borne sensors may provide an additional way to obtain these observations. Biologging devices have become progressively smaller and can now collect environmental information together with animal movement data (Kays et al. 2015; Wilmers et al. 2015). Flying birds are particularly interesting in this context because they naturally move across different parts of the urban landscape and at different altitudes. Previous studies have demonstrated the potential of bird-borne sensors for atmospheric monitoring. Thomas et al. (2018) developed and tested miniature meteorological sensors carried by eagles, obtaining vertical air-temperature profiles. In Rome, Di Bernardino et al. (2022) used homing pigeons equipped with GPS-linked atmospheric sensors to investigate spatial variation in CO₂ revealing differing CO₂ concentrations decreasing from central urban areas towards the periphery and lower concentrations over urban green spaces. Although air temperature was also recorded, the analysis primarily focused on CO₂ and did not reconstruct the three-dimensional thermal structure of the urban atmosphere. The integration of temperature observations from species with complementary flight characteristics to reconstruct the three-dimensional thermal structure of an entire urban area remains largely unexplored.

Here, we used GPS-linked air-temperature measurements collected by homing pigeons (*Columba livia*) and yellow-legged gulls (*Larus michahellis*) to describe the spatial, vertical and daytime structure of the lower urban atmosphere over Rome, Italy. We combined bird-borne observations with information on altitude above ground level, distance from the city centre, land cover, tree-patch area and topography in a spatial modelling framework. We addressed three main questions: (1) how is the urban landscape associated with local variation in air temperature; (2) how does the centre-to-periphery air-temperature gradient change with altitude and time of day; and (3) how do the vertical extent and spatial organization of warm atmospheric layers change across the city and during the day? Finally, we used the model predictions to reconstruct and visualize the atmospheric thermal field across the study area.

## 2. Materials and methods

### 2.1 Study area and data collection

The study was conducted in Rome, central Italy, within a 17-km radius from a reference point in the historical city centre (12.4902° E, 41.8905° N). The study area included densely built-up areas, urban green spaces, semi-natural areas and peripheral sectors of the city, covering a broad centre-to-periphery gradient and a wide range of ground elevations.

Air temperature was measured using bird-borne sensors carried by homing pigeons and yellow-legged gulls. The final dataset included 47 individuals: 42 pigeons and five yellow-legged gulls. Pigeon data were collected during summer sampling campaigns in 2021, 2023, 2024 and 2025, whereas gull data were collected between June and August 2024. The five gulls were adults captured at their nests and equipped with backpack-mounted devices secured with Teflon threads. All five birds were released immediately after device attachment.

Data were recorded using Axy-Trek data loggers (TechnoSmart Europe, Italy) equipped with an external fast-response temperature sensor. The temperature probe was exposed to ambient air and positioned away from the bird’s body to reduce interference from body heat. The sensor response time was ≤ 1 s, and the same temperature-sensor configuration was used for both species. Pigeon GPS and temperature data were recorded at 1 Hz, while gull measurements were recorded at 5-min intervals.

### 2.2 Data preparation

Spatial coordinates were projected to WGS 84 / UTM zone 32N (EPSG:32632). Distance from the reference point in the historical city centre was calculated for each observation and expressed in kilometres. The projected coordinates were also used in the two-dimensional spatial smooth included in the statistical models.

Air temperature (°C) was used as the response variable. Explanatory variables included hour of day, altitude above ground level (AGL), distance from the city centre, ground elevation, land-cover class and tree-patch area. Altitude AGL was calculated by subtracting local ground elevation from the altitude above sea level recorded by the bird-borne logger.

Land-cover information was obtained from a supervised classification of Sentinel-2 imagery acquired during summer 2024 at 10 × 10 m spatial resolution (European Space Agency; Drusch et al. 2012). Classification was performed in QGIS using the Semi-Automatic Classification Plugin (Congedo 2021). The resulting raster was used to assign a land-cover class to each bird observation and prediction-grid cell. Four classes were initially considered: artificial surfaces, semi-natural non-woody vegetation, semi-natural woody vegetation and water. No valid observations classified as water remained after filtering, so the final analyses included the first three classes. The 2024 classification was used for all sampling years as a common representation of land cover within the study area.

Ground elevation was obtained from the Digital Elevation Model (DEM) described by Tarquini and Nannipieri (2017) and extracted for both bird observations and prediction-grid cells. Ground elevation was used to calculate altitude AGL and was also included as a model covariate. Tree-patch area was derived from the land-cover classification. For observations and prediction-grid cells classified as semi-natural woody vegetation, the area of the corresponding woody patch was calculated and assigned to the point. Patch area was expressed in hectares for analysis.

Analyses were restricted to observations collected during June, July and August and between 07:00 and 18:00 local time, focusing on the daytime summer thermal structure. The dataset was screened for outliers (Andersen 2008; Bjørneraas et al. 2010), and only non-stationary observations below 400 m AGL and within 17 km of the central reference point were retained. Observations containing missing values for temperature, hour, altitude, ground elevation, land cover or distance from the city centre were excluded.

A spatial prediction grid with a resolution of 100 × 100 m was prepared across the 17-km study domain and linked to the environmental variables required for prediction.

Before model fitting, descriptive analyses summarized the range and spatial coverage of the observations. Summary statistics were calculated for air temperature, altitude AGL, distance from the city centre, ground elevation and tree-patch area, while sampling effort was summarized by bird species and land-cover class (Tables S1–S2). Air-temperature and altitude distributions were examined separately for pigeons and gulls, and observations were mapped to assess geographic coverage.

### 2.3 Statistical modelling of the urban thermal field

Air temperature was modelled using generalized additive models (GAMs), fitted with the bam() function of the mgcv package in R. Models assumed Gaussian errors and an identity link. The analysis was designed to describe the centre-to-periphery and vertical temperature gradients while accounting for temporal variation, environmental variables and residual spatial structure.

Model selection was carried out in two stages. First, four candidate models were compared to determine the functional form of the centre-to-periphery and vertical temperature gradients. All models included a smooth effect of hour of day (k = 9), land-cover class, tree-patch area, ground elevation, a distance-by-altitude interaction, a two-dimensional thin-plate regression spline of projected coordinates (k = 40), and sampling date as a random-effect smooth. The spatial smooth accounted for geographic variation not captured by distance from the city centre or the environmental variables. The four models differed in the form assigned to distance and altitude: linear distance and linear altitude (M1), linear distance and quadratic altitude (M2), quadratic distance and linear altitude (M3), and quadratic distance and quadratic altitude (M4).

The best-supported model from this first comparison was then extended to test whether the horizontal and vertical temperature gradients changed during the day. A distance-by-hour interaction was added first (M2b), followed by an altitude-by-hour interaction (M2c). The final M2c structure therefore included a linear centre-to-periphery gradient, a quadratic altitude effect and interactions between distance and altitude, distance and hour, and altitude and hour (Table S3).

All candidate models were fitted using maximum likelihood (ML) and compared using Akaike’s Information Criterion (AIC). Models were ranked by AIC, and ΔAIC was calculated relative to the model with the lowest value. The best-supported model was then refitted using fast restricted maximum likelihood (fREML) for parameter estimation, inference and prediction. The complete structure of the final model is reported in Table S3.

Model adequacy was evaluated using standard GAM residual diagnostics and checks of basis-dimension adequacy. Concurvity was also examined to assess overlap among model terms. Residual spatial autocorrelation was assessed using Moran’s I on residuals aggregated to 1-km spatial cells, using eight nearest neighbours and 999 permutations.

Predictive performance was evaluated using an internal 80:20 hold-out validation stratified by sampling date. Within each date, observations were randomly assigned to a training dataset (80%) and a test dataset (20%), so that all sampling dates were represented in both datasets. The final model structure was fitted to the training data and used to predict air temperature for the withheld observations. Prediction accuracy was assessed using root mean square error (RMSE), mean absolute error (MAE) and the coefficient of determination (R²) between observed and predicted temperatures (Willmott and Matsuura 2005; Chai and Draxler 2014).

Spatial predictions were generated from the final model across the 17-km study domain. The random effect of sampling date was excluded so that the resulting maps represented the general thermal structure rather than conditions associated with a specific sampling date. All other model terms, including the two-dimensional spatial smooth, were retained. Air temperature was predicted at 09:00, 13:00 and 17:00 and at 20, 150 and 300 m AGL. These altitudes represented lower, intermediate and upper parts of the sampled atmospheric range while avoiding predictions close to the sparsely sampled upper limit.

### 2.4 Three-dimensional reconstruction of the urban thermal field

The three-dimensional thermal field was reconstructed by extending the horizontal prediction grid from 0 to 400 m AGL at 20-m intervals. Air temperature was predicted at each altitude and selected hour using the final GAM and the same prediction settings described above.

Isothermal surfaces of 34, 35 and 36 °C were reconstructed at 13:00 to describe the vertical extent of warm air across the study area. For each horizontal grid cell, the predicted vertical temperature profile was examined to identify the height at which air temperature crossed each selected threshold. Crossing height was estimated by linear interpolation between consecutive altitude levels, allowing isotherm height to vary continuously despite the 20-m spacing of the prediction grid.

The primary isothermal boundary was defined as the height at which temperature decreased from above to below the selected threshold with increasing altitude. It therefore represented the upper limit of the atmospheric layer warmer than that temperature. Because the final model included a quadratic altitude effect, some predicted vertical profiles were non-monotonic and crossed the same threshold more than once. In these cases, the warm-to-cool crossing was retained as the primary boundary, while subsequent crossings in which temperature increased again above the threshold were recorded separately as secondary crossings. Their frequency and vertical position were summarized in Table S9.

To examine centre-to-periphery variation, the heights of the 34, 35 and 36 °C isotherms at 13:00 were grouped into 1-km distance bands from the historical city centre. Median height and interquartile range were calculated within each band for each temperature threshold.

Daytime variation was examined using the 34 °C isotherm, which was reconstructed at 09:00, 13:00 and 17:00 using the same procedure. At each hour, its vertical position was summarized by the median height and interquartile range (IQR). Spatial occurrence was calculated as the proportion of horizontal prediction-grid cells in which a primary crossing occurred within the 0–400 m AGL range. These measures were used to compare changes in the height and spatial distribution of the 34 °C isotherm during the day.

## 3. Results

### 3.1 Sampling coverage and descriptive characteristics

The final dataset included 61,917 air-temperature observations from 47 individuals across 91 sampling dates (Table S1). Pigeons contributed 59,585 observations from 42 individuals (96.2% of the dataset), while yellow-legged gulls contributed 2,332 observations from five individuals (3.8%). Artificial surfaces accounted for 32,401 observations (52.3%), semi-natural non-woody vegetation for 23,767 (38.4%) and semi-natural woody vegetation for 5,749 (9.3%). Observations extended from the historical city centre to approximately 17 km distance, with a mean distance of 9.22 ± 3.35 km (Fig. 1a). Air temperature ranged from 17.1 to 42.9 °C, with a mean of 29.03 ± 2.96 °C. Altitude ranged from near-ground conditions to 399.98 m AGL, although most observations were concentrated at lower altitudes (median = 51.1 m AGL; IQR = 30.0–93.8 m; Fig. S1A,B; Table S2). Gull observations occurred at higher altitudes than pigeon observations, with median altitudes of 73.4 and 50.6 m AGL, respectively (Figs. 1B and S1C). Sampling covered the full retained daytime period from 07:00 to 18:00, with the greatest sampling effort around midday (Fig. S1D).

**Fig. 1.**
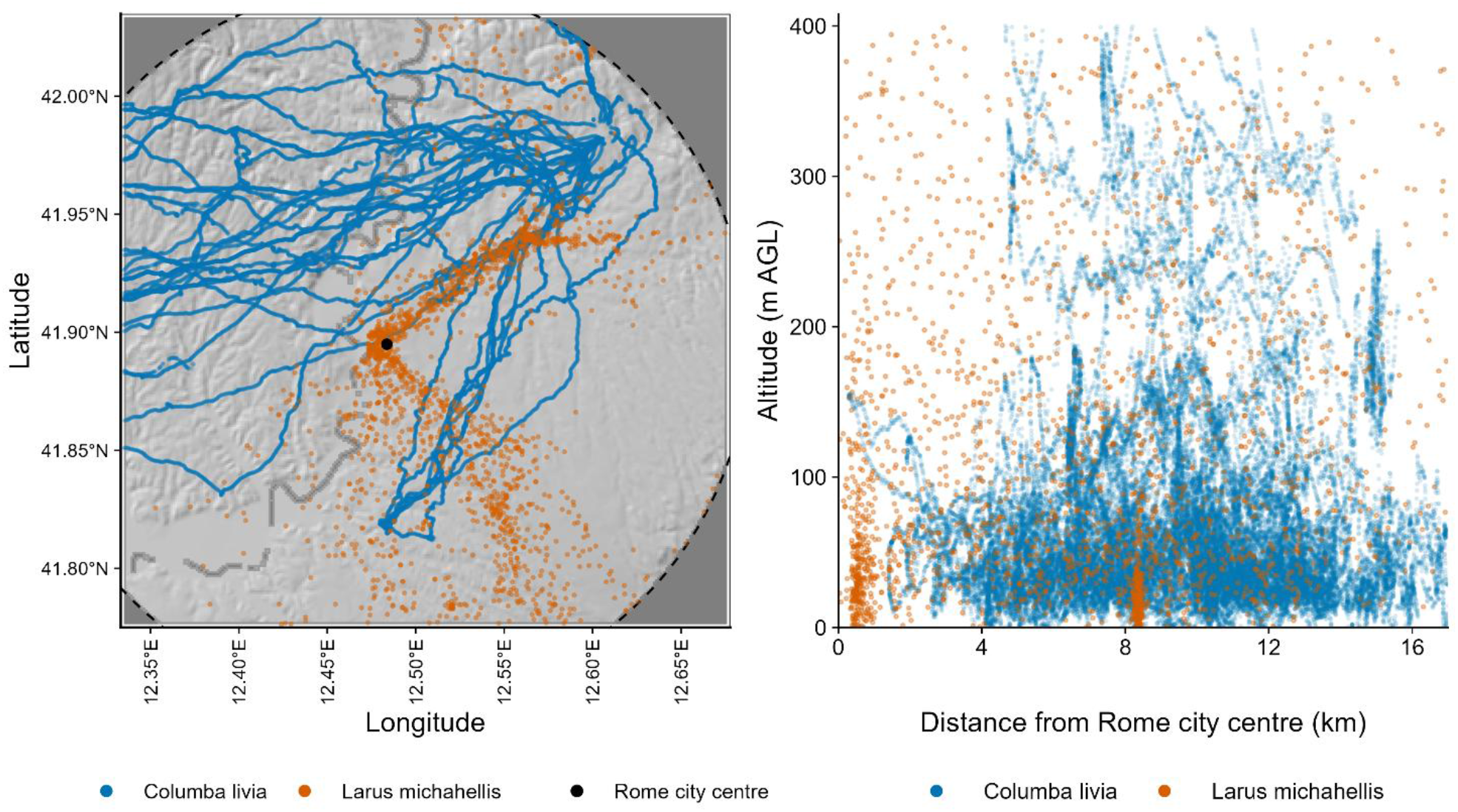
Spatial and vertical coverage of bird-borne air-temperature observations across the study area. (a) Geographic distribution of observations collected by homing pigeons and yellow-legged gulls within the 17-km study domain. The black point indicates the reference location in the historical city centre. (b) Distribution of observations across distance from the city centre and altitude above ground level (AGL), shown separately for the two species.

### 3.2 Model selection, model fit and predictive performance

Model comparison supported a linear centre-to-periphery temperature gradient combined with a quadratic altitude effect. Among the four initial candidate models, M2 had the lowest AIC, outperforming models with either a quadratic distance term or a linear altitude effect (Table 1). Adding the distance-by-hour interaction further improved model support, followed by an additional improvement after including the altitude-by-hour interaction. The resulting M2c model had the lowest AIC (222,242.00), with ΔAIC = 112.97 relative to the next-best model, M2b, and was therefore selected as the final model.

**Table 1.** Comparison of candidate generalized additive models describing the spatial, vertical and daytime structure of urban air temperature. Models are ranked according to Akaike’s Information Criterion (AIC), with ΔAIC calculated relative to the best-supported model. D and A denote distance from the city centre and altitude above ground level, respectively.

| Model | Distance | Altitude | Additional temporal interactions | df | AIC | $\Delta AIC$ |
| --- | --- | --- | --- | --- | --- | --- |
| M2c | Linear | Quadratic | $D \times \text{Hour} + A \times \text{Hour}$ | 147.27 | 222242 | 0 |
| M2b | Linear | Quadratic | $D \times \text{Hour}$ | 146.17 | 222355 | 112.97 |
| M2 | Linear | Quadratic | — | 145.09 | 223115 | 873.04 |
| M4 | Quadratic | Quadratic | — | 146.07 | 223200 | 957.61 |
| M1 | Linear | Linear | — | 144.13 | 223851 | 1609.11 |
| M3 | Quadratic | Linear | — | 144.94 | 223862 | 1620.09 |

The final model explained 75.80% of the deviance in air temperature and had an adjusted R² of 0.76. It included the distance-by-altitude interaction specified a priori, together with distance-by-hour and altitude-by-hour interactions (Table S5). Ground elevation was negatively associated with air temperature (β = −0.00495 °C m⁻¹, p < 0.001). Compared with artificial surfaces, semi-natural non-woody vegetation had lower predicted temperatures (β = −0.0962 °C, p < 0.001), whereas semi-natural woody vegetation did not differ significantly (β = −0.0140 °C, p = 0.558). Tree-patch area was also negatively associated with air temperature (β = −0.000690 °C ha⁻¹, p = 0.006).

Residual diagnostics showed some deviation from Gaussianity in the tails of the residual distribution (Fig. S2). Observed concurvity was 0.49 for the hour smooth, 0.64 for the two-dimensional spatial smooth and 0.89 for the sampling-date random effect (Table S4A). Basis-dimension diagnostics indicated remaining fine-scale structure in the spatial smooth, which was not substantially reduced by increasing the basis dimension (Table S4B). Spatially aggregated residuals retained positive spatial autocorrelation (Moran’s I = 0.22, p = 0.001; 999 permutations; Fig. S3).

In the internal hold-out validation, the model fitted to the 80% training subset predicted the withheld observations with an RMSE of 1.50 °C, an MAE of 1.03 °C and an R² of 0.75.

### 3.3 Three-dimensional predictions of the urban thermal field

#### 3.3.1 Horizontal and vertical thermal structure

Model predictions showed a clear centre-to-periphery temperature gradient that varied with altitude and time of day (Fig. 2; Table S6). Air temperature decreased with increasing distance from the city centre, but the gradient weakened with altitude. At 09:00, it decreased from −0.40 °C km⁻¹ at 20 m AGL (95% CI = −0.51 to −0.29) to −0.22 °C km⁻¹ at 300 m AGL (95% CI = −0.33 to −0.11). At 13:00, the corresponding gradient decreased from −0.30 °C km⁻¹ (95% CI = −0.41 to −0.18) to −0.12 °C km⁻¹ (95% CI = −0.23 to −0.01). By 17:00, it had weakened further, from −0.19 °C km⁻¹ at 20 m AGL (95% CI = −0.30 to −0.08) to −0.02 °C km⁻¹ at 300 m AGL (95% CI = −0.13 to 0.10). At this time, confidence intervals included zero at both 150 and 300 m AGL.

**Fig. 2.**
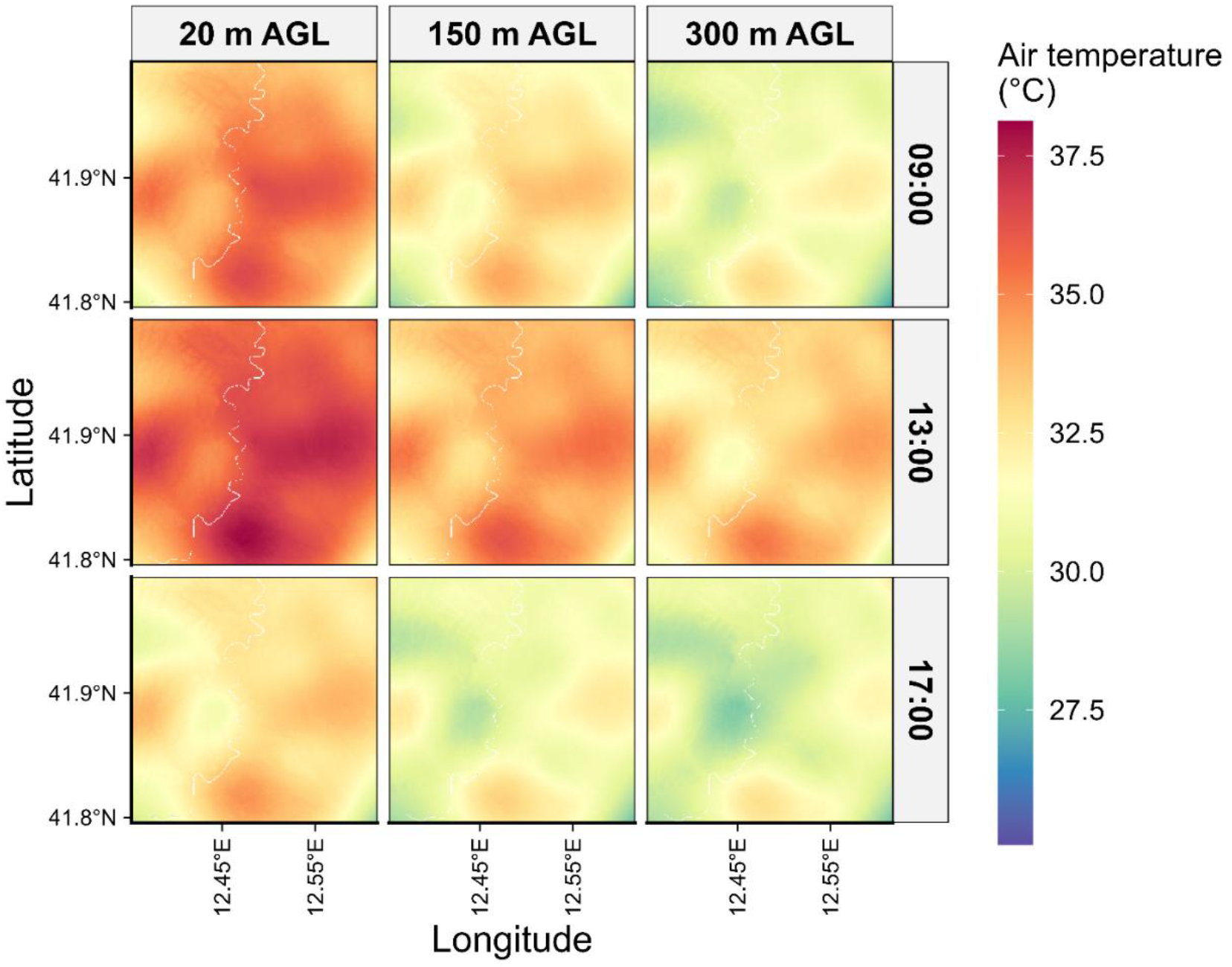
Predicted air temperature across the study domain at different hours and altitudes. Model predictions are shown at 09:00, 13:00 and 17:00 (rows) and at 20, 150 and 300 m above ground level (AGL; columns). Predictions were obtained from the final GAM excluding the sampling-date random effect. A common temperature scale is used across all panels.

Within this broad centre-to-periphery pattern, horizontal predictions also showed local spatial variation across the study domain (Fig. 2). This variation was most evident at lower altitude, particularly around midday, and decreased at 150 and 300 m AGL.

#### 3.3.2 Isothermal geometry at 13:00

At 13:00, isotherm height decreased as the temperature threshold increased (Fig. 3; Fig. S4; Online Resource 2). The 34 °C isotherm had a median height of 135.0 m AGL (IQR = 88.8–172.0 m), compared with 76.1 m AGL for the 35 °C isotherm (IQR = 42.7–110.0 m) and 37.1 m AGL for the 36 °C isotherm (IQR = 17.4–66.5 m). Maximum reconstructed heights were 358, 330 and 182 m AGL for the 34, 35 and 36 °C isotherms, respectively. Secondary crossings occurred in 10.8% of grid cells with a primary 34 °C crossing and 2.15% of those with a primary 35 °C crossing, while none occurred for the 36 °C isotherm (Table S9A) Median isotherm height generally decreased with distance from the city centre for all three temperature thresholds (Fig. 4). The 36 °C isotherm was more spatially restricted than the 34 and 35 °C isotherms, occurring in 34,598 grid cells compared with 55,632 and 56,212 cells, respectively (Table S7). Where a primary 36 °C crossing occurred, its height ranged from near-ground level to 182 m AGL (Table S7), with the highest values confined to localized areas of the study domain (Fig. 5).

**Fig. 3.**
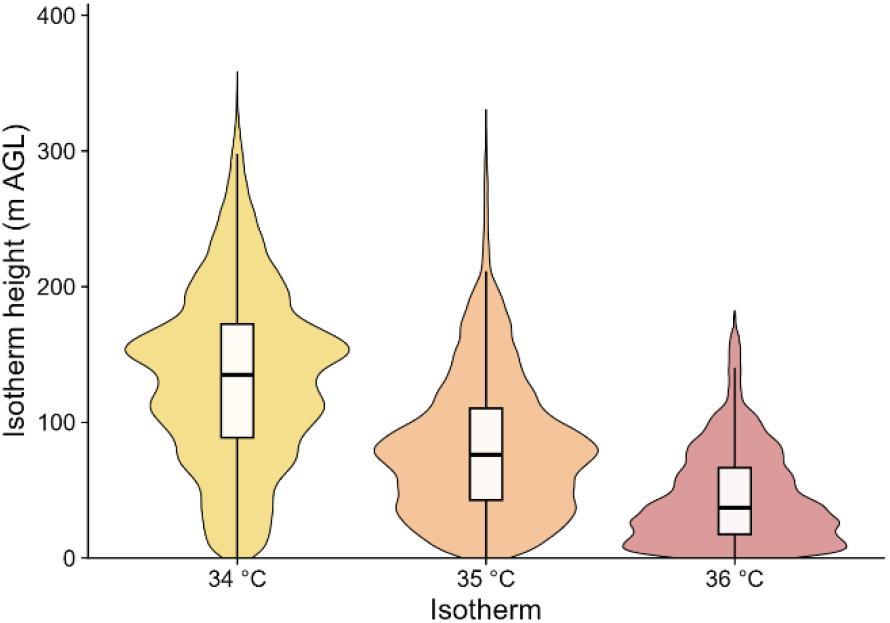
Distribution of primary isotherm heights above ground level (AGL) for the three temperature thresholds. Boxplots show the median and interquartile range within each distribution.

**Fig. 4.**
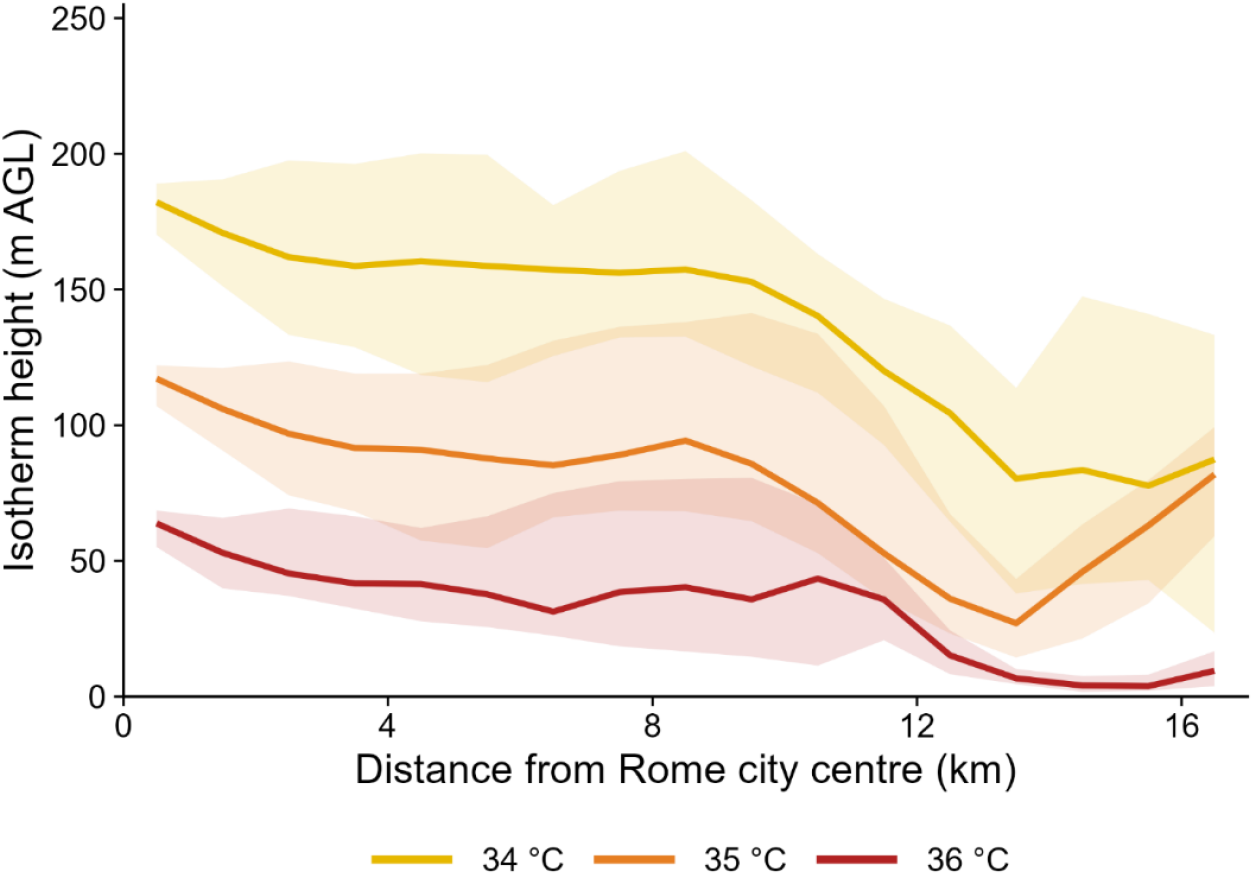
Centre-to-periphery variation in reconstructed isotherm height at 13:00. Median heights above ground level (AGL) of the primary 34, 35 and 36 °C isothermal boundaries are shown across 1-km distance bands from the historical city centre. Shaded areas represent the interquartile range within each distance band.

**Fig. 5.**
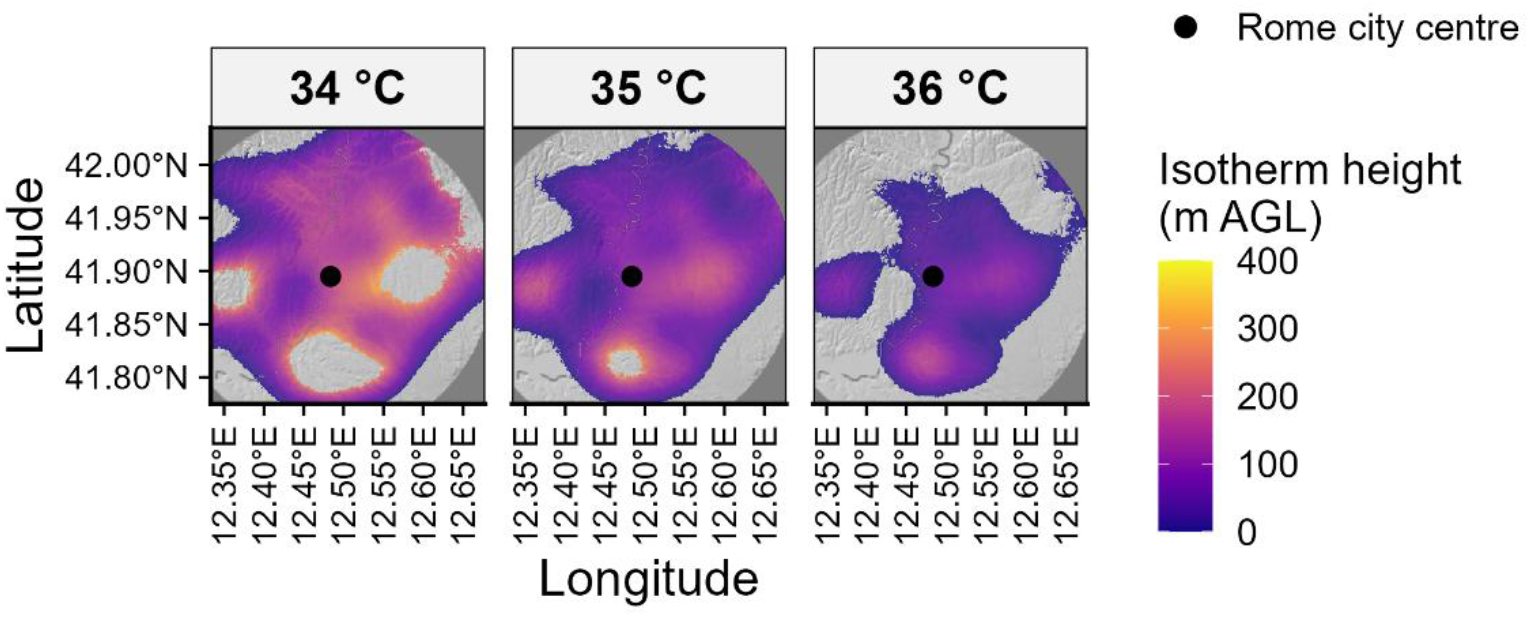
Spatial distribution of reconstructed isotherm heights at 13:00. Maps show the height above ground level (AGL) of the primary 34, 35 and 36 °C isothermal boundaries across the study domain. Grey areas indicate grid cells in which no primary warm-to-cool crossing was reconstructed within the 0–400 m AGL prediction range. The black point indicates the historical city centre.

#### 3.3.3 Daytime evolution of the 34 °C isotherm

The height of the 34 °C isotherm changed markedly during the day (Fig. 6A; Table S8). At 09:00, its median height was 56.8 m AGL (IQR = 28.5–89.6 m), with a maximum reconstructed height of 185 m. At 13:00, the median increased to 135.0 m AGL (IQR = 88.8–172.0 m) and the maximum to 358 m. By 17:00, the median height had decreased to 14.0 m AGL (IQR = 6.3–31.8 m), with a maximum of 82 m (Fig. 6B).

**Fig. 6.**
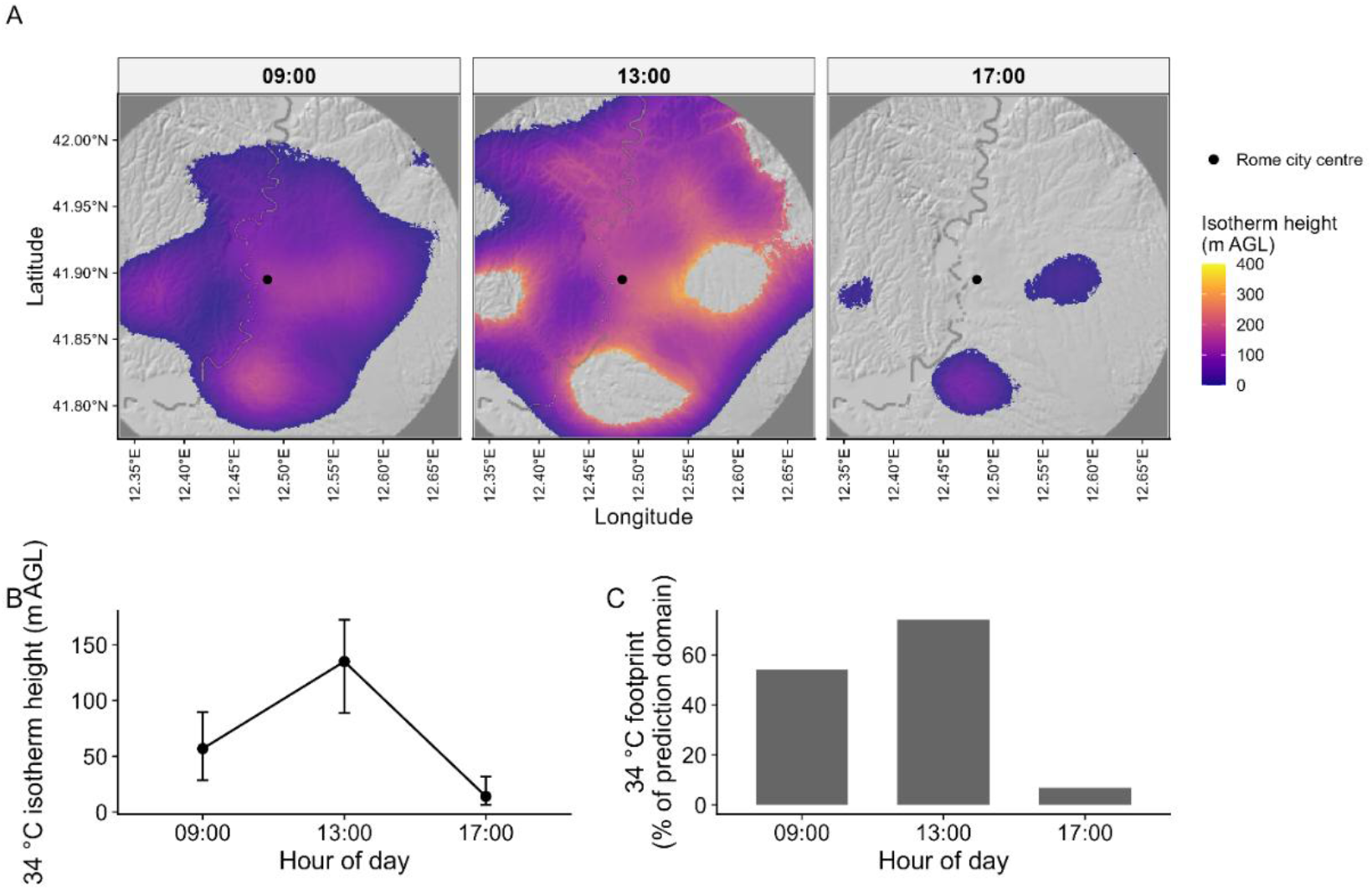
Daytime variation in the reconstructed 34 °C isothermal boundary. (a) Spatial distribution of primary 34 °C isotherm height above ground level (AGL) at 09:00, 13:00 and 17:00. Grey areas indicate grid cells without a primary warm-to-cool crossing within the 0–400 m AGL prediction range. (b) Median isotherm height at each hour; error bars represent the interquartile range. (c) Proportion of the prediction domain in which a primary 34 °C crossing was reconstructed.

Spatial occurrence followed the same daytime pattern. Primary 34 °C crossings occurred in 54.0% of the prediction domain at 09:00, increased to 74.0% at 13:00 and decreased to 6.9% at 17:00 (Fig. 6C).

The 34 °C boundary therefore reached its greatest vertical extent and spatial occurrence at 13:00 and was much lower and less widespread by 17:00.

## 4. Discussion

Our results show that the urban atmospheric thermal field over Rome had a complex spatial structure extending from the surface into the lower atmosphere. Local temperature variation was associated with characteristics of the underlying landscape, including land cover, tree-patch area and ground elevation. This local spatial heterogeneity was embedded within a broader thermal structure that varied with distance from the city centre, altitude and time of day. Air temperature generally decreased from the city centre towards the periphery, but this gradient weakened with altitude and through the daytime period. At the same time, the vertical extent of high-temperature air changed substantially during the day and reached its greatest development around midday. These patterns show that horizontal thermal contrast and the vertical extent of warm air represent related but distinct features of the urban thermal environment.

### 4.1 Urban landscape and local air-temperature variation

The spatial heterogeneity in air temperature across Rome is consistent with the heterogeneous nature of the urban landscape. Different land-cover types can modify surface energy exchanges through differences in radiation absorption, heat storage, water availability and evapotranspiration. Differences in air temperature among urban land uses have been reported in several cities (Shudo et al. 1997), while tree cover and urban green spaces can contribute to local cooling through shading and evapotranspiration (Shashua-Bar and Hoffman 2000). The magnitude of these effects, however, varies considerably among sites, vegetation types and climatic conditions (Bowler et al. 2010). Studies in Lisbon and Hong Kong, for example, reported spatially variable cooling associated with urban green spaces (Oliveira et al. 2011; Ng et al. 2012). In our study, semi-natural non-woody areas were slightly cooler than artificial surfaces. For woody vegetation, the model separated the contribution of land-cover class from that of tree-patch area: the broad woody land-cover class showed little difference from artificial surfaces, whereas larger tree patches were associated with slightly lower air temperatures. These results suggest that broad land-cover classes alone provide only a partial description of local variation in urban air temperature.

The spatial extent of woody vegetation may therefore be important in addition to its categorical presence. Tree-patch size can influence local thermal conditions through changes in shading and transpiration (Jiao et al. 2017), while studies at broader spatial scales have shown that vegetation– temperature relationships can also depend on landscape configuration and the surrounding urban matrix (Guo et al. 2020; Jung et al. 2021). Zhou et al. (2022) further showed that the relative importance of landscape characteristics can change with the spatial scale considered. Much of this literature is based on land surface temperature and therefore cannot be directly compared with our measurements of air temperature. Nevertheless, it shows that a simple woody/non-woody classification may mask important differences among vegetated areas. The limited difference associated with the broad woody land-cover class, together with the negative association with tree-patch area, is consistent with the possibility that patch extent contains information that is not captured by land-cover class alone. The magnitude of the tree-patch association was small, however, and should not be interpreted as a direct estimate of the cooling produced by individual patches.

This distinction is also relevant to the type of measurements used here. In Rome, Gratani and Varone (2006) reported lower air temperatures associated with urban trees, while Gratani et al. (2016) measured local microclimatic differences related to vegetation within an urban park. These studies mainly characterized conditions beneath or close to tree canopies, where the influence of shading and evapotranspiration is most direct. Our observations instead covered a broad range of heights above ground. The influence of immediate surface conditions may therefore be less directly represented in bird-borne measurements than in measurements made within or close to the canopy. The relatively weak land-cover contrasts observed here are thus not necessarily inconsistent with stronger local cooling reported beneath trees or within parks. Because land-cover and tree-patch effects were estimated across the sampled altitude range, they should also be interpreted as average associations rather than as height-specific cooling effects.

Ground elevation represented an additional component of spatial variation. Topography can modify urban temperature patterns together with land use, vegetation and urban morphology (Tzavali et al. 2015). The negative association observed in Rome shows that terrain-related variation needs to be considered when interpreting intra-urban temperature differences. Local landscape characteristics therefore contributed to the heterogeneity of the atmospheric thermal field, but they represented only one component of its spatial structure. This local variation was embedded within a broader temperature field that also changed systematically with distance from the city centre, altitude and time of day.

### 4.2 Centre-to-periphery and vertical structure of the urban thermal field

A broad centre-to-periphery temperature gradient was evident across the study area, although the predicted thermal field did not show a uniform or concentric decrease in temperature from the city centre (Fig. 2). Several warmer and cooler areas occurred across the domain, indicating substantial spatial heterogeneity superimposed on the overall radial tendency. Despite this heterogeneity, the model-derived centre-to-periphery gradient showed a clear overall pattern: its magnitude decreased both with increasing altitude and through the daytime period (Table S6).

The vertical weakening of the gradient was evident at all three times considered. At 09:00, for example, its magnitude decreased from −0.397 °C km⁻¹ at 20 m AGL to −0.220 °C km⁻¹ at 300 m AGL. The same pattern occurred at 13:00 and 17:00, and by 17:00 the gradient was no longer clearly distinguishable from zero at 150 and 300 m AGL. Thus, although local thermal heterogeneity remained evident across the city, the broad centre-to-periphery organization became progressively less pronounced with height. Similar height-dependent thermal contrasts have been reported in previous urban observations. Duckworth and Sandberg (1954), using simultaneous vertical temperature profiles over built-up and peripheral open areas, found that near-surface temperature differences decreased with height and that the profiles could occasionally cross. Saaroni et al. (2000) also found that the magnitude of the urban thermal contrast differed between street and roof levels in Tel Aviv. More recently, three-dimensional modelling over Bologna showed a progressive attenuation of the urban–rural temperature contrast with height within the lower atmosphere (Possega et al. 2026). These studies differ in spatial scale, observational design and definition of the urban thermal contrast, so direct quantitative comparisons with Rome are not appropriate (Tzavali et al. 2015).

The radial gradient also weakened through the daytime period. At 20 m AGL, its magnitude decreased from −0.397 °C km⁻¹ at 09:00 to −0.295 °C km⁻¹ at 13:00 and −0.193 °C km⁻¹ at 17:00, and the same general temporal pattern occurred at higher altitudes (Table S6). The broad centre-to-periphery thermal contrast was therefore strongest in the morning and became progressively less pronounced through the day. This does not necessarily indicate faster warming of the city centre. Rather, it shows that temperatures across the study domain became less strongly differentiated according to distance from the centre as the day progressed. Strong daytime variability in urban thermal contrasts has also been reported in other cities. In Beijing, Yang et al. (2013) observed a marked spatial organization from the inner urban belts towards the outer parts of the city, but the strength of the urban thermal contrast changed considerably during the day and was relatively weak during the warm daytime hours. Studies in Mediterranean cities also show that urban temperature contrasts vary strongly with time of day and meteorological conditions (Giannaros and Melas 2012; Papanastasiou and Kittas 2012). The temporal weakening observed over Rome should therefore be interpreted as a change in the strength of the broad radial component of the thermal field, rather than as a uniform daytime change in temperature across the city.

The reconstruction of isothermal boundaries provided a complementary view of the vertical organization of this thermal field. At 13:00, progressively higher temperature thresholds occurred at progressively lower altitudes: the median height decreased from about 135 m AGL for the 34 °C boundary to 76 m for 35 °C and 37 m for 36 °C (Table S7). Isotherm height also showed an overall decrease from the city centre towards the periphery (Fig. 4), but the reconstructed surfaces were spatially heterogeneous rather than smooth and concentric (Figs. 3–5). This organization shares some qualitative features with the idealized concept of an urban heat dome, in which the thermally affected atmospheric layer is deeper over warmer urban areas (Hidalgo et al. 2008; Fan et al. 2017). Real urban thermal fields, however, are often more complex. Kim and Baik (2005), for example, reported multiple warm cores within the Seoul UHI, while observations in Lisbon showed substantial spatial variability associated with urban position, topography and local and regional wind systems (Lopes et al. 2013). Numerical studies further show that background flow can displace and deform idealized urban thermal structures (Zhang et al. 2025). The heterogeneous geometry reconstructed over Rome is therefore compatible with the complexity expected in a real urban atmosphere. However, fine-scale spatial features should be interpreted cautiously because observational coverage was not uniform across the prediction domain.

The 34 °C boundary also changed markedly through the daytime period (Fig. 6; Table S8). Its median height increased from about 57 m AGL at 09:00 to 135 m at 13:00, before decreasing to about 14 m at 17:00. Its occurrence across the prediction domain followed the same broad pattern, being greatest at 13:00 and strongly reduced by 17:00. The upward displacement of this boundary indicates that air at or above 34 °C occupied a deeper portion of the lower atmosphere around midday. Interestingly, this maximum vertical development did not coincide with the strongest centre-to-periphery gradient, which occurred at 09:00. The two patterns therefore describe complementary aspects of the thermal field: the radial gradient describes its broad horizontal organization, whereas the isothermal boundaries describe the vertical extent of high-temperature air. Thus, while the centre-to-periphery contrast weakened from morning to midday, the high-temperature layer simultaneously expanded vertically. One possible interpretation is that daytime heating affected an increasingly deep atmospheric layer while horizontal temperature differences became less pronounced. Urban boundary-layer studies show that daytime heating and mixing can modify both the horizontal and vertical thermal structure of cities (Miao et al. 2009; Hidalgo et al. 2010; Pal et al. 2012). However, turbulence, mixed-layer depth and advection were not measured here, so these processes remain possible mechanisms rather than directly observed causes.

This combination of horizontal and vertical changes shows that the daytime thermal structure over Rome cannot be described by a single measure of urban temperature contrast. The reconstructed isotherms shared some geometric features with the conceptual urban heat dome, but their heterogeneous shape and temporal variation also illustrate the complexity of a real urban atmosphere. Temperature measurements alone cannot establish the circulation responsible for such a structure, which would require information on wind, convergence and vertical motion (Fan et al. 2017). Nevertheless, the combined analysis of radial gradients and isothermal surfaces reveals an organized three-dimensional thermal field in which a broad centre-to-periphery tendency, local spatial heterogeneity and the vertical development of high-temperature air changed together through the daytime period.

### 4.3 Limitations and methodological considerations

Some limitations should be considered when interpreting these results. First, the dataset was strongly unbalanced between species, with only five gulls contributing a relatively small proportion of the observations. Although gulls extended the spatial and vertical sampling coverage, the reconstructed thermal field was primarily informed by pigeon observations. Nevertheless, observational coverage remained uneven across the prediction domain, and some residual spatial structure persisted. Fine-scale features of the predicted maps and isothermal surfaces should therefore be interpreted more cautiously than the broader patterns consistently identified across the domain.

Second, the study describes the thermal structure of the lower urban atmosphere but does not directly resolve the atmospheric processes producing it. Wind, atmospheric stability, turbulence and boundary-layer depth were not measured simultaneously with temperature, although these factors are known to influence urban thermal contrasts and their spatial organization (Morris et al. 2001; Mihalakakou et al. 2002; Lee and Baik 2010). Vertical mixing, advection and urban-breeze circulation therefore remain possible interpretations rather than mechanisms directly demonstrated by our data.

Finally, the reconstructed fields represent average summer daytime patterns across multiple sampling dates rather than individual meteorological episodes. Differences among 09:00, 13:00 and 17:00 should therefore be interpreted as systematic hour-related changes in the fitted thermal structure rather than as the evolution of a single day. Future studies combining animal-borne temperature sensing with simultaneous meteorological observations and more spatially balanced sampling could further resolve the processes shaping the three-dimensional urban thermal field.

## 5. Conclusions

This study shows that the urban atmospheric thermal field over Rome was organized across both horizontal and vertical dimensions. A broad centre-to-periphery temperature gradient was superimposed on substantial spatial heterogeneity, and its magnitude decreased both with increasing altitude and through the daytime period. The reconstruction of isothermal surfaces provided a complementary view of this structure, showing that high-temperature air extended well above the urban surface and that its vertical and spatial organization changed markedly during the day. In particular, the strongest centre-to-periphery contrast occurred in the morning, whereas the 34 °C boundary reached its greatest vertical extent around midday. These results show that horizontal thermal contrast and the vertical development of high-temperature air represent distinct but complementary properties of the urban thermal field.

Beyond these climatological results, the study demonstrates the potential of animal-borne sensing for observing atmospheric conditions at spatial and vertical scales that are difficult to sample using conventional monitoring networks. To our knowledge, this is the first study to combine temperature observations from two complementary bird species to reconstruct the three-dimensional thermal structure of the atmosphere across an entire urban area. Pigeons provided dense observations of the lower urban atmosphere, while the higher and wider-ranging movements of gulls extended sampling to atmospheric layers and parts of the city that would otherwise be more difficult to observe. Combining such mobile observations with spatial modelling therefore offers a complementary approach to fixed meteorological stations, remote sensing and atmospheric modelling. More broadly, integrating animal-borne sensing with conventional meteorological observations could provide new opportunities to characterize the vertical dimension and daytime dynamics of urban heat.

## Supplementary Information

Online Resource 1. Supplementary tables (S1–S9) and figures (S1–S4) reporting sampling characteristics, model specifications, diagnostic analyses and additional results supporting the main manuscript.

Online Resource 2. Rotating three-dimensional visualization of the reconstructed primary 34, 35 and 36 °C isothermal boundaries over Rome at 13:00, showing their spatial variation and vertical position relative to local ground elevation.

## Declarations

### Data availability

The data supporting the findings of this study are available from the corresponding author upon reasonable request.

### Author contributions

Giacomo Grosso: Conceptualization, Data curation, Formal analysis, Investigation, Methodology, Writing – original draft. Claudio Carere: Supervision. Giacomo Dell’Omo: Resources, Supervision, Writing – review and editing. All authors read and approved the final manuscript.

### Use of generative AI in manuscript preparation

During the preparation of this work the authors used ChatGPT (OpenAI) and Claude (Anthropic) to improve language quality, structure sentences, and assist in drafting some parts of the manuscript.

## Supporting information

all supplementary non video material

## Acknowledgments

We gratefully acknowledge the support of the *Ornis italica* Association for logistical assistance and access to the homing pigeon facilities used during the study.

