## Supplementary material for "Beyond the urban surface: bird-borne sensing reveals the atmospheric structure of urban heat": all supplementary non video material

**Table S1. Sampling characteristics of the bird-borne temperature dataset used in the final analysis.** The table reports the number of individuals, observations, sampling dates, sampling period and retained hourly range for each bird species. The total number of sampling dates represents unique dates across the complete dataset; some dates included observations from both species.

| **Table S1. Sampling characteristics of the bird-borne temperature dataset used in the final analysis.** | | | | | |
|---|---|---|---|---|---|
| Species | Individuals (n) | Observations (n) | Sampling dates (n) | Sampling period | Hour range |
| Yellow-legged gulls (G) | 5 | 2,332 | 85 | 1 Jun–31 Aug 2024 | 07:00–18:00 |
| Feral pigeons (P) | 42 | 59,585 | 11 | 3 Jul 2021–18 Jun 2025 | 07:00–18:00 |
| Total | 47 | 61,917 | 91* | 2021–2025 | 07:00–18:00 |

**Table S2. Descriptive characteristics and sampling composition of the final modelling dataset.** Summary statistics are reported for air temperature, altitude above ground level (AGL), distance from the city centre, ground elevation and tree-patch area, together with the number and percentage of observations by species and land-cover class. ART = artificial surfaces; SNL = semi-natural woody vegetation; SNNL = semi-natural non-woody vegetation.

| **Table S2 variables** | | | | | | | | | |
|---|---|---|---|---|---|---|---|---|---|
| Variable | Mean ± SD | Median | Q25–Q75 | Range | | Variable | Category | n | % |
| Temperature (°C) | 29.03 ± 2.96 | 29.1 | 27.10 – 30.50 | 17.10–42.90 | | Species | Yellow-legged gull | 2,332 | 3.77 |

| | | | | | | | | | |
|---|---|---|---|---|---|---|---|---|---|
| Altitude (m AGL) | 77.10 ± 73.28 | 51.14 | 29.99–93.78 | <0.01–399.98 | | | Feral pigeon | 59,585 | 96.23 |
| Distance from city centre (km) | 9.22 ± 3.35 | 9.42 | 6.72–11.62 | 0.03–17.00 | | Land cover | (ART) | 32,401 | 52.33 |
| Ground elevation (m a.s.l.) | 58.45 ± 30.34 | 50.71 | 36.90–74.04 | 5.08–251.54 | | | (SNL) | 5,749 | 9.29 |
| Tree-patch area (ha) | 4.05 ± 28.15 | 0 | 0.00–0.00 | 0.00–307.63 | | | (SNNL) | 23,767 | 38.39 |

**Table S3. Structure of the candidate generalized additive models used to describe the urban atmospheric thermal field.** Candidate models differed in the functional form of distance from the city centre and altitude and in the inclusion of distance-by-hour and altitude-by-hour interactions. All models included a distance-by-altitude interaction, a smooth effect of hour, land-cover class, tree-patch area, ground elevation, a two-dimensional spatial smooth and sampling date as a random-effect smooth.

| Table S3. Structure of candidate generalized additive models evaluated for reconstructing the three-dimensional urban atmospheric thermal field. | | | | | | | |
|---|---|---|---|---|---|---|---|
| Model | Distance from city centre | Altitude | Distance × Altitude | Distance × Hour | Altitude × Hour | Spatial smooth | Date effect |
| M1 | Linear | Linear | ✓ | – | – | ✓ | ✓ |
| M2 | Linear | Quadratic | ✓ | – | – | ✓ | ✓ |
| M2b | Linear | Quadratic | ✓ | ✓ | – | ✓ | ✓ |
| M2c | Linear | Quadratic | ✓ | ✓ | ✓ | ✓ | ✓ |
| M3 | Quadratic | Linear | ✓ | – | – | ✓ | ✓ |

| | | | | | | | |
|---|---|---|---|---|---|---|---|
| M4 | Quadratic | Quadratic | ✓ | − | − | ✓ | ✓ |

**Table S4A. Concurvity diagnostics for the final generalized additive model.** Worst, observed and estimate concurvity measures are reported for each model component. Values closer to 1 indicate greater overlap among model terms. **B. Basis-dimension diagnostics for smooth terms in the final generalized additive model.** The table reports the available basis dimension (k′), estimated degrees of freedom (edf), k-index and associated p-value for each smooth term. A k-index is not reported for the sampling-date random-effect smooth.

| **Table S4A — Concurvity** | | | |
|---|---|---|---|
| Model component | Worst | Observed | Estimate |
| Parametric component | 1 | 1 | 1 |
| Hour, (s(Hour)) | 0.952 | 0.487 | 0.861 |
| Spatial smooth, (s(x,y)) | 0.999 | 0.638 | 0.4 |
| Sampling date, (s(Date)) | 1 | 0.89 | 0.417 |

| **Table S4B — Basis-dimension (k) diagnostics of the final GAM** | | | | |
|---|---|---|---|---|
| Smooth term | k′ | edf | k-index | p-value |
| s(Hour) | 8 | 7.91 | 0.979 | 0.022 |
| s(x_km,y_km) | 39 | 38.54 | 0.496 | <0.001 |
| s(Date) | 91 | 88.3 | — | — |

**Table S5. Parametric coefficients and smooth terms of the final generalized additive model for air temperature.** Parametric terms are reported with coefficient estimates, standard errors (SE), *t* statistics and p-values; smooth terms are reported with estimated degrees of freedom (edf), reference degrees of freedom (Ref.df), *F* statistics and p-values. Artificial surfaces were used as the reference land-cover class. Because distance, altitude and hour occur in interaction terms, their individual coefficients are conditional on the values of the interacting variables.

| **Table S5 — Final model** | | | | |
|---|---|---|---|---|
| **Term** | **Estimate** | **SE** | **t** | **p** |
| Intercept | 37.38 | 0.6482 | 57.668 | <0.001 |
| Land cover: semi-natural woody | −0.0140 | 0.024 | −0.585 | 0.558 |
| Land cover: semi-natural non-woody | −0.0962 | 0.0137 | −7.029 | <0.001 |
| Tree-patch area (ha) | −0.000690 | 0.000251 | −2.749 | 0.006 |
| Ground elevation (m a.s.l.) | −0.00495 | 0.00048 | −10.321 | <0.001 |
| Distance from city centre (km) | −0.6395 | 0.0574 | −11.146 | <0.001 |
| Altitude (m AGL) | −0.03251 | 0.000825 | −39.408 | <0.001 |
| $\text{Altitude}^2$ | $2.907 \times 10^{-5}$ | $1.002 \times 10^{-6}$ | 28.996 | <0.001 |
| Distance × Altitude | $6.306 \times 10^{-4}$ | $3.136 \times 10^{-5}$ | 20.106 | <0.001 |
| Distance × Hour | 0.02553 | 0.000879 | 29.062 | <0.001 |
| Altitude × Hour | $6.218 \times 10^{-4}$ | $5.101 \times 10^{-5}$ | 12.191 | <0.001 |
| **Smooth term** | **edf** | **Ref.df** | **F** | **p** |
| Hour of day | 7.91 | 8 | 25,970.6 | <0.001 |
| Spatial smooth, s(x, y) | 38.54 | 39 | 1,455.7 | <0.001 |
| Sampling date | 88.3 | 90 | 846.3 | <0.001 |

**Table S6. Vertical position of reconstructed isothermal boundaries at 13:00.** The table summarizes the number of prediction-grid cells containing a primary warm-to-cool crossing and the distribution of reconstructed heights above ground level (AGL) for the 34, 35 and 36 °C isotherms.

| **Table S6 — Isotherms at 13:00** | | | | | |
|---|---|---|---|---|---|
| Isotherm | Grid cells (n) | Mean ± SD (m AGL) | Median (m AGL) | Q25–Q75 (m AGL) | Range (m AGL) |
| 34 °C | 55,632 | 133.0 ± 64.2 | 135 | 88.8–172.0 | 0.01–358.0 |
| 35 °C | 56,212 | 82.2 ± 51.6 | 76.1 | 42.7–110.0 | <0.01–330.0 |
| 36 °C | 34,598 | 45.1 ± 34.7 | 37.1 | 17.4–66.5 | <0.01–182.0 |

**Table S7. Daytime variation in the reconstructed 34 °C isothermal boundary.** The table reports the number and proportion of prediction-grid cells containing a primary warm-to-cool crossing and summary statistics for isotherm height above ground level (AGL) at 09:00, 13:00 and 17:00.

| **Table S7 — Daytime evolution of the 34 °C isotherm** | | | | | |
|---|---|---|---|---|---|
| Time | Grid cells with primary crossing (n) | Prediction domain (%) | Mean ± SD (m AGL) | Median (m AGL) | Q25–Q75 (m AGL) |
| 09:00 | 40,649 | 54% | 61.2 ± 39.8 | 56.8 | 28.5–89.6 |
| 13:00 | 55,632 | 74% | 133.0 ± 64.2 | 135 | 88.8–172.0 |
| 17:00 | 5,168 | 6.87% | 21.3 ± 19.8 | 14 | 6.3–31.8 |

**Table S8. Model-derived centre-to-periphery air-temperature gradients at selected hours and altitudes.** Gradients are reported for 09:00, 13:00 and 17:00 at 20, 150 and 300 m above ground level (AGL), together with standard errors and 95% confidence intervals. Negative values indicate decreasing air temperature with increasing distance from the city centre. Values represent the model-derived radial gradient conditional on the specified hour and altitude and do not include local variation represented by the two-dimensional spatial smooth.

| **Table S8. Model-derived centre-to-periphery temperature gradients at selected hours and altitudes.** | | | | |
|---|---|---|---|---|
| Time | Altitude (m AGL) | Gradient (°C $km^{-1}$) | SE | 95% CI |
| 09:00 | 20 | −0.397 | 0.057 | −0.508 to −0.286 |
| 09:00 | 150 | −0.315 | 0.057 | −0.426 to −0.204 |
| 09:00 | 300 | −0.220 | 0.057 | −0.332 to −0.109 |
| 13:00 | 20 | −0.295 | 0.057 | −0.406 to −0.184 |
| 13:00 | 150 | −0.213 | 0.057 | −0.324 to −0.102 |
| 13:00 | 300 | −0.118 | 0.057 | −0.230 to −0.007 |
| 17:00 | 20 | −0.193 | 0.057 | −0.304 to −0.081 |
| 17:00 | 150 | −0.111 | 0.057 | −0.222 to 0.001 |
| 17:00 | 300 | −0.016 | 0.057 | −0.128 to 0.096 |

**Table S9A. Frequency and vertical position of primary and secondary isothermal crossings at 13:00.** Primary crossings represent warm-to-cool transitions and define the upper boundary of the layer exceeding each temperature threshold. Secondary crossings represent subsequent cool-to-warm transitions at higher altitude. The table reports their frequency and median height with interquartile range for the 34, 35 and 36 °C thresholds. **B. Frequency and vertical position of primary and secondary 34 °C isothermal crossings during the daytime period.** Primary and secondary crossings are summarized at 09:00, 13:00 and 17:00. Primary crossings represent warm-to-cool transitions, whereas secondary crossings represent subsequent cool-to-warm transitions at higher altitude. Heights are reported as median and interquartile range above ground level (AGL).

| **Table S9A. Frequency and vertical position of primary and secondary isothermal crossings at 13:00.** | | | | | |
|---|---|---|---|---|---|
| Isotherm | Primary crossings (n) | Cells with two crossings (n) | Two crossings (%) | Primary height, median [IQR] (m AGL) | Secondary height, median [IQR] (m AGL) |
| 34 °C | 55,632 | 6,010 | 10.8 | 135.1 [88.8–172.5] | 362.9 [333.9–383.3] |
| 35 °C | 56,212 | 1,208 | 2.15 | 76.1 [42.7–110.3] | 379.5 [363.4–389.5] |
| 36 °C | 34,598 | 0 | 0 | 37.1 [17.4–66.5] | — |
| **Table S9B — 34 °C during daytime** | | | | | |
| Time | Primary crossings (n) | Cells with two crossings (n) | Two crossings (%) | Primary height, median | Secondary height, median [IQR] (m AGL) |

| | | | | | |
|---|---|---|---|---|---|
| | | | | [IQR] (m AGL) | |
| 09:00 | 40,649 | 0 | 0 | 56.8 [28.5–89.6] | — |
| 13:00 | 55,632 | 6,010 | 10.8 | 135.1 [88.8–172.5] | 362.9 [333.9–383.3] |
| 17:00 | 5,168 | 3 | 0.058 | 14.0 [6.3–31.8] | 396.1 [392.8–398.3] |

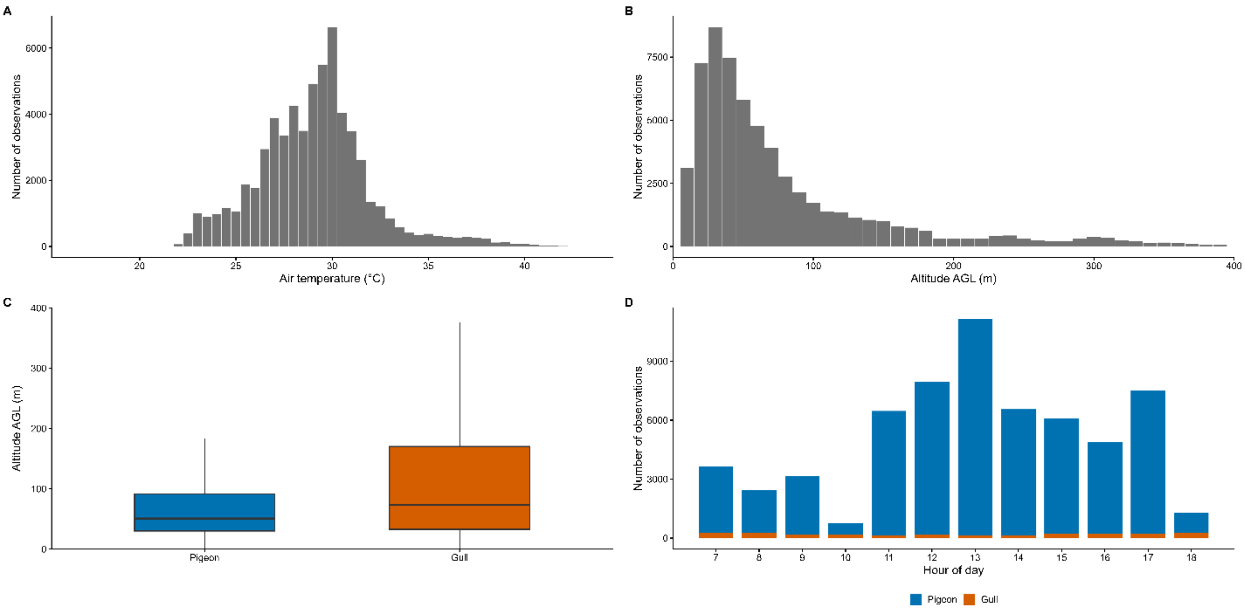

**Figure S1. Descriptive characteristics and sampling distribution of the final bird-borne temperature dataset.** (A) Distribution of air-temperature observations. (B) Distribution of observations across altitude above ground level (AGL). (C) Altitude distribution for feral pigeons and yellow-legged gulls. (D) Number of observations collected during each hour of the retained daytime period, shown separately for the two species.

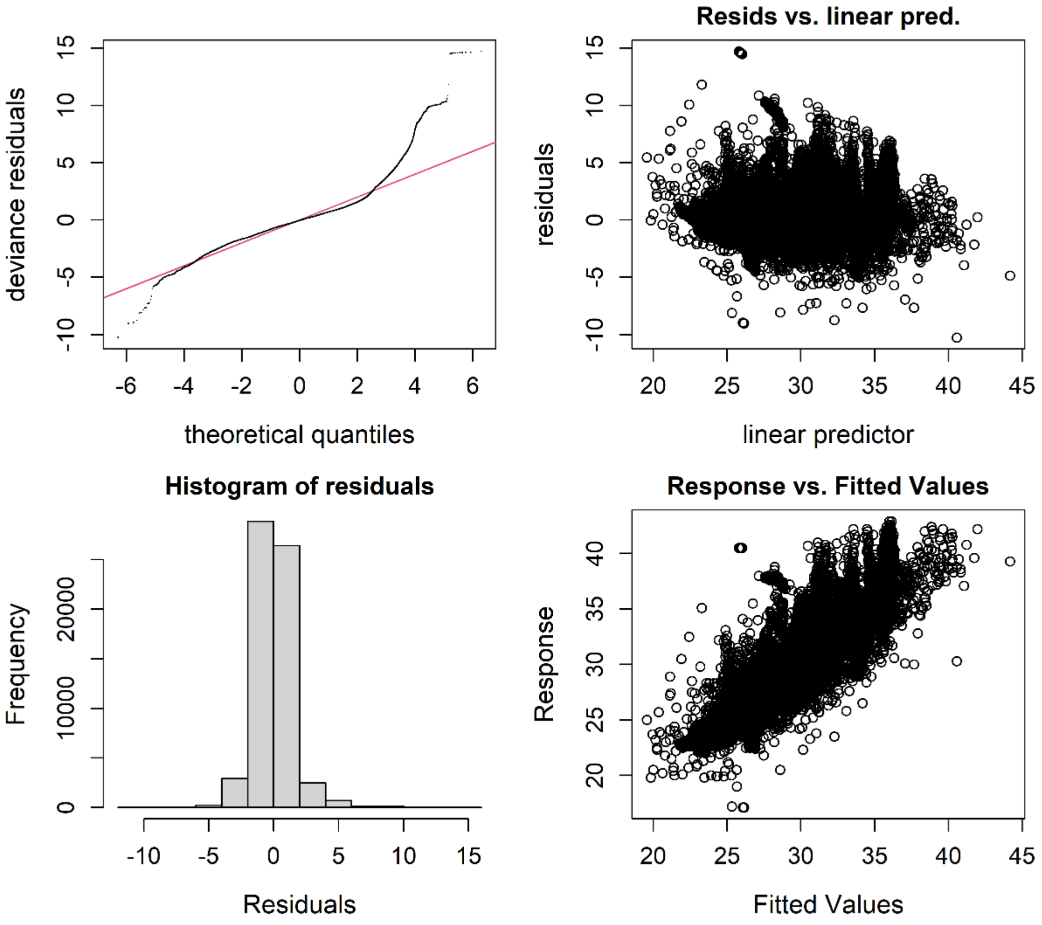

**Figure S2. Diagnostic plots for the final generalized additive model.** Panels show the quantile–quantile plot of deviance residuals, residuals against the linear predictor, the residual-frequency distribution, and observed air temperature against fitted values.

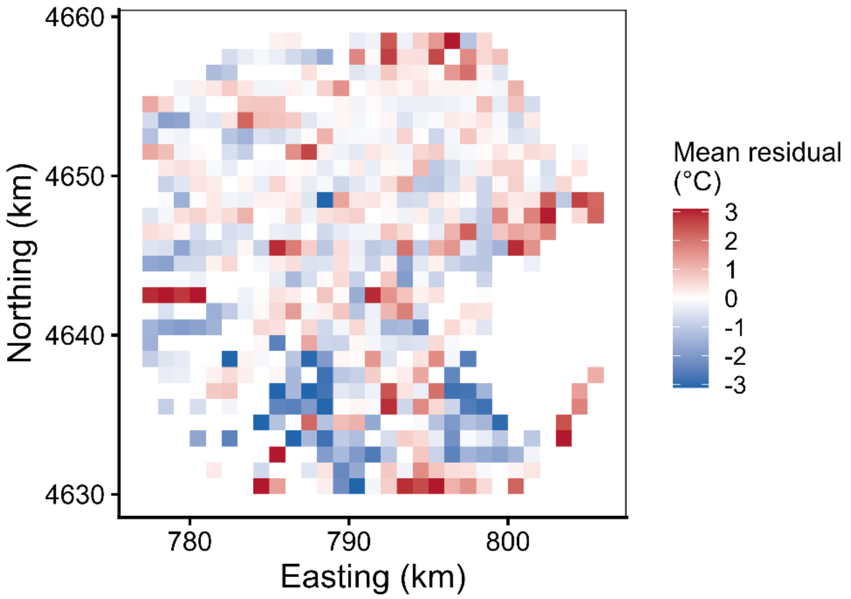

**Figure S3. Spatial distribution of residuals from the final generalized additive model.** Residuals were aggregated as mean values within 1-km spatial cells across the study domain. Positive and negative values indicate areas where observed temperatures were respectively higher or lower than model predictions.

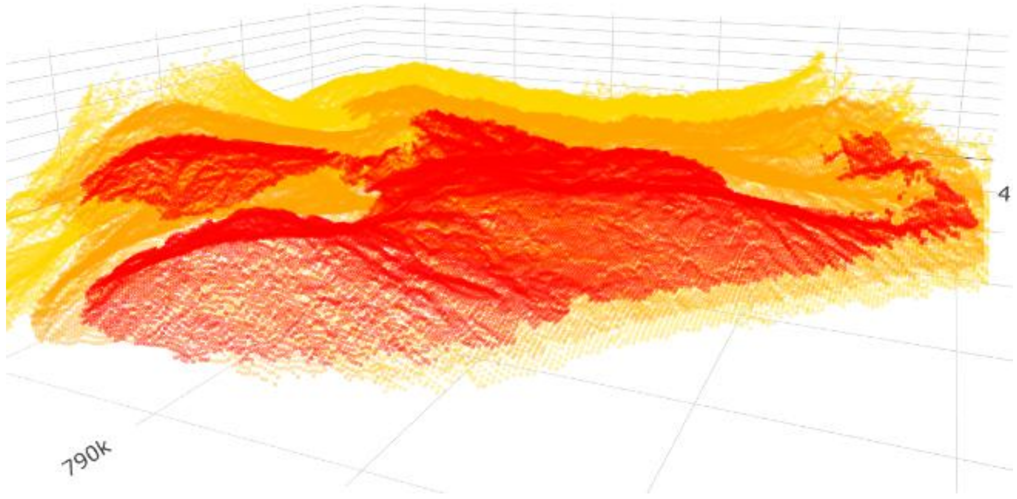

Figure S4. Three-dimensional structure of the reconstructed isothermal boundaries at 13:00. Reconstructed 34, 35 and 36 °C primary isothermal boundaries across the study domain. The three-dimensional visualization combines isotherm height above ground level with local ground elevation.
